# Mechanical and thermodynamic properties of *Aβ*_42_, *Aβ*_40_, and *α*-synuclein fibrils: A coarse-grained method to complement experimental studies

**DOI:** 10.1101/527242

**Authors:** Adolfo B. Poma, Horacio V. Guzman, Mai Suan Li, Panagiotis E. Theodorakis

**Affiliations:** Institute of Fundamental Technological Research, Polish Academy of Sciences, Paw-ińskiego 5B, 02-106 Warsaw, Poland; Max Planck Institute for Polymer Research, Ackermannweg 10, 55128 Mainz, Germany; Institute of Physics, Polish Academy of Sciences, Al. Lotników 32/46, 02-668 Warsaw, Poland

**Keywords:** Atomic Force Microscopy, *β*-amyloid, *α*-synuclein, mechanical deformation, molecular simulation, proteins

## Abstract

We perform molecular dynamics simulation on several relevant biological fibrils associated with neurodegenerative diseases such as *Aβ*_40_, *Aβ*_42_, and *α*-synuclein systems to obtain a molecular understanding and interpretation of nanomechanical characterization experiments. The computational method is versatile and addresses a new subarea within the mechanical characterization of heterogeneous soft materials. We investigate both the elastic and thermodynamic properties of the biological fibrils in order to substantiate experimental nanomechanical characterization techniques that are quickly developing and reaching dynamic imaging with video rate capabilities. The computational method qualitatively reproduces results of experiments with biological fibrils, validating its use in extrapolation to macroscopic material properties. Our computational techniques can be used for the co-design of new experiments aiming to unveil nanomechanical properties of biological fibrils from a molecular understanding point of view. Our approach allows a comparison of diverse elastic properties based on different deformation, *i.e*. tensile (*Y*_L_), shear (*S*), and indentation (*Y*_T_). From our analysis, we find a significant elastic anisotropy between axial and transverse directions (*i.e. Y*_T_ > *Y*_L_) for all systems. Interestingly, our results indicate a higher mechanostability in the case of *Aβ*_42_ fibrils than in the case of *Aβ*_40_, suggesting a significant correlation between mechanical stability and aggregation propensity (rate) in amyloid systems, that is, the higher the mechanical stability the faster the fibril formation. Finally, we find that α-synuclein fibrils are thermally less stable than *β*-amyloid fibrils. We anticipate that our molecular-level analysis of the mechanical response under different deformation conditions for the range of fibrils considered here will provide significant insights for the experimental observations.

**Background:** Nanomechanical characterization of a single biological fibril is generally a challenge due to the typical thermal motion. Here, we propose a computational protocol that can assist experiment in elucidating the molecular background of the mechanical response in fibrils related to neurodegenerative diseases.

**Results:** We performed a systematic comparison of mechanical properties of different biological fibrils involved in neurodegenerative diseases. Our results show a higher mechanocanostability in case of *Aβ*_42_ fibrils than in the case of *Aβ*_40_. This effect is observed for all different types of mechanical deformation. Moreover, the *α*-synuclein fibril shows a large anisotropy (i.e. *Y*_T_ > *Y*_L_) in comparison with *β*-amyloid fibrils, and it is thermally less stable than *β*-amyloid fibrils.

## Introduction

All-atom molecular dynamics (MD) simulation has been employed to study the physical and chemical behaviour of the fundamental biomolecules of life (*e.g*. proteins [1], nucleic acids [2] and lipids [3]). To this end, lipid membranes, viral capsids, and biological fibrils are common examples of large complexes that pose significant challenges for all-atom simulation. For example, the time scales of various biological processes are in the range of 10^−6^ − 10^−3^ s, and thus they are orders of magnitude larger than typical molecular motion (i.e. 10^−15^ − 10^−12^ s) captured in all-atom MD. The length scales are similarly much smaller in all-atom simulation than it would be relevant for studying processes involving large conformation changes in large biological complexes. In the context of mechanical properties of various fibrils, for example, *β*-amyloids [4,5], cellulose [6] and collagen [7], all-atom models have been used to estimate the elastic moduli based on the response of the system, but mostly approximately. Still, molecular-level methods are necessary to understand the microscopic mechanisms of the mechanical response of biological fibrils. In this regard, coarse-grained (CG) models are suitable, because they remove several degrees of freedom of the system, which enables them to reach the experimental time and length scales that describe the relevant phenomena while maintaining a molecular-level description of the systems under consideration [8–11]. In particular, CG simulation is able to describe large structural changes in the context of fibril deformation, which would be otherwise impossible with all-atom models. In particular, the CG model can be used to infer the elastic parameter in ideal conditions, which is given by the Hertz model [12] and is valid for isotropic materials and as close as possible to the experimental conditions [13]. While other sophisticated ‘Hertz models’ [14,15] aim to study the elastic properties of anisotropic materials with high symmetries, *e.g*. crystals, softer materials such as biological fibrils or polymers are not suitable for such descriptions. Although biological matter is an example of an anisotropic material, it is not expected to follow *a priori* a simple Hertzian relationship given by *F* ≈ *Y_T_h*^3/2^ (with *Y_T_* the transversal Young modulus and *h* the indentation depth). When it actually follows this relationship, the elastic modulus can be easily obtained from the slope of the curve. This approach can be used to test the experimental estimation of an elastic property. Most importantly, the mechanism of deformation that give rise to the linear response can be characterized in the CG simulation. From the experimental point of view, there is a long-standing discussion in the Atomic Force Microscopy (AFM) community whether Hertzian mechanics is applicable to all explored soft matter samples with AFM. One of the basic assumptions of Hertz model is that the indented object is a half-space and made out of a homogeneous material. However, at the nanoscale it is intrinsically difficult to measure pure and homogeneous materials, or perfectly mixed materials, with some exceptional cases, such as the Highly Oriented Pyrolytic Graphite (HOPG), Silica, and other ‘clean’ surfaces, which are, however, very far away from biological systems. Moreover, by considering the indenter as a sphere, the anisotropies in the deformed material can be screened, since the measured deformation depends on the contact area, which will be the arc region that forms in contact with the sphere. In considering other shapes for the cantilever tip, such as conical or flat punch, the impact of the anisotropy is expected to be much higher [16]. Nonetheless, to our knowledge the exact shape of the cantilever tip cannot be determined during experimental measurements. As a result, big discrepancies are found when comparing Young moduli measured with macroscopic techniques and nanoscopic ones such as AFM, because a nanoscopic exploration of biological systems reaches molecular resolutions and the measurements are in general very delicate due to the intrinsic properties of soft matter and the danger of damaging the samples [17]. As a matter of fact, the employed reference model to study the mechanical response of the biological fibrils during AFM nanoindentation has been also the Hertz model. Hence we also use it as a reference for comparing the indentational values we obtained to the experimental ones, although we remark that our molecular modeling can adapt further anisotropic mechanical models, envisioned within force microscopy techniques.

Biological fibrils are well known biomaterials of practical use. The related technological applications range from drug delivery [18] to structural scaffolds [19], where the role of the fibril may be to immobilize small molecules (*e.g*. enzymes [20]). The applications are motivated by their unique properties, such as the spontaneous formation under certain conditions, the high mechanical stability (comparable to silk), and the ability of forming ordered structures, albeit the monomeric units (proteins) of these fibrils are intrinsically disordered. [21,22] These are fundamental properties for applications that require that fragmentation of the material be avoided, for example, during synthesis, active process (drug delivery) or response to an external perturbation (*e.g*. change in temperature). To this end, the interplay between mechanical and thermodynamic properties will determine the overall behaviour of the fibrils, which depends on the arrangement of the individual amino acid chains in these structures. The case of fibrils consisting of either 40-mer or 42-mer amyloid chains (it contains two additional hydrophobic amino acids) is particularly interesting. For example, A*β*_40_ typically assembles into two-fold and three-fold symmetries (see Fig. 1), while the highest symmetry reported by experiments for A*β*_42_ fibrils is a two-fold symmetry, as in the case of *α*-synuclein (*α*-syn) fibrils. [23,24] Furthermore, the aggregation typically takes place 2.5 times faster in a solution of *Aβ*_42_ than in the case of *Aβ*_40_ [25,26]. Interestingly, the aggregation rate of fibril formation has been found to be highly correlated with the mechanical properties of the fibrils, namely, the mechanically more stable fibril is the one with faster aggregation [27]. While experimental observations have been derived from a small set of samples, our CG simulations can be used to validate these observations and study a larger set of fibrils.

**Figure 1:**
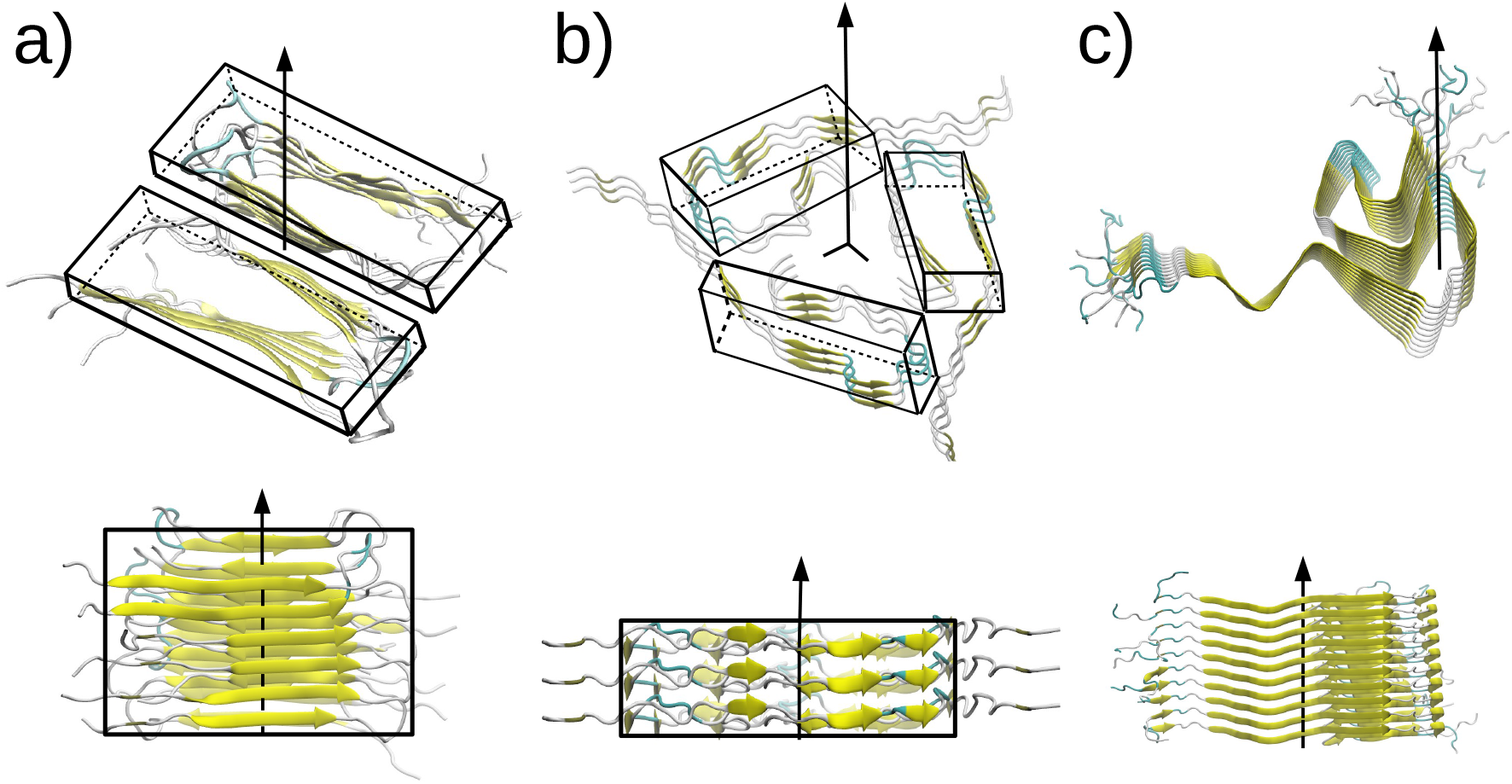
Snapshots illustrate a part of biological fibrils used in our simulation. The main axis of symmetry is indicated and the secondary structure for each chain. Panel (a) illustrates a *β* A_40_ (PDB id: 2LMO) with two-fold symmetry, while panel (b) a *β* A_40_ fibril (PDB id: 2M4J) with three-fold symmetry. Panel (c) illustrates the *α*-syn fibril (PDB id: 2N0A) with no symmetry. Rectangular boxes depict the local symmetry.

Typical length scales of biological fibrils are in the range between nm and *μ*m, therefore, AFM, which can operate, for example, in static (contact) and dynamic modes, has been one of the main methods to study such systems [28,29]. On the one hand, AFM in contact-mode has been used to provoke the mechanical deformation of fibrils, in this way obtaining the Young modulus (here denoted as *Y*_T_) [30–32]. On the other hand, the experimental determination of the tensile Young’s modulus (*Y*_L_) is nontrivial at the nanoscale [33], due to the requirement of a different experimental setup, namely, the more involved sonification method [34]. Moreover, the experimental calculation of the shear modulus (*S*) can be realised by suspending the fibril between two beams and pressing the free part against the indenter, which gives rise to the fibril bending modulus (*Y*_b_) that depends on both the *Y*_T_ and the *S*.

In this respect, our CG strategy can be used to extract and compare elastic properties in a systematic way. This significant advantage of CG simulation has motivated the current study, which employs MD simulation of a structure-based CG model [35–38] to investigate one *α*-synuclein and five *β*-amyloid fibrils of known experimental structure related to specific neurodegenerative diseases. Our simulation sheds light on the mechanical and thermodynamic properties of these fibrils by providing the microscopic picture required to explain the relevant phenomena. We achieve this by applying different types of deformation (*e.g*. tension, shearing, indentation) and analysing the intermolecular contacts between amino acids. Our simulations reveal significant differences in the mechanical behaviour between 40 and 42 *β*-amyloid, and *α*-syn fibrils. Moreover, we find that the *α*-syn fibril is thermally less stable than the *β*-amyloid fibrils.

In the next section, we present details about our methodology. Then, we present our results and analysis, and in the last section we summarise our conclusions.

## Materials and Methods

To realise our studies, we have chosen three different A*β*_40_ fibrils with PDB ids: 2LMO[39], 2M4J[40] and 2MVX[41] and two A*β*_42_ with PDB ids: 5OQV[42], and 2NAO[43]. The only available structure for *α*-syn is the one with PDB id: 2N0A[44].

### The coarse-grained model

In our CG model, each amino acid is represented by a bead located at the *C_α_*-atom position. The potential energy between beads reads:

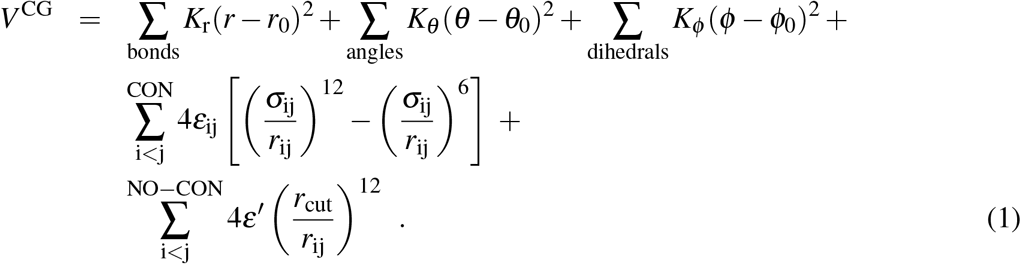

The first three terms on the right hand side of Eq.(1) correspond to the harmonic pseudo-bond, bond angle and dihedral potentials. The values of the elastic constants is, *K_r_* = 100 kcal/mol/Å^2^, *K_θ_* = 45 kcal/mol/rad^2^ and *K_ϕ_* = 5.0 kcal/mol/rad^2^, which were derived from all-atom simulation[45]. The choice of equilibrium values r_0_, *θ*_0_, and *ϕ*_0_ are based on two, three, and four α-C atoms, respectively, and are meant to favour the native geometry. The fourth term on the right-hand side of Eq.(1) takes into account the non-bonded contact interactions, described by the Lennard–Jones (LJ) potential. Here, we take *ε*_ij_ to be uniform and equal to *ε* = 1.5 kcal/mol, which is also derived by all-atom simulation [45]. Our approach has shown very good agreement with experimental data on stretching [46,47] and nanoindentation of biological fibrils, such as virus capsids [35] and *β*-amyloids [36]. The strength of the repulsive non-native term, *ε*′, is set equal to *ε*. Our CG model takes into account native distances as in the case of a Gō-like model[37]. Hence, the native contacts are determined by the overlap criterion [48]. In practice, each heavy atom is assigned to a van der Waals radius, as proposed by Tsai *et al*. [49]. A sphere with the radius enlarged by a factor of 1.24 is built around the atom. If two amino acids have heavy atoms with overlapping spheres, then we consider a native contact between those two *C_α_* atoms. In Fig. 2, we show the CG representation for some biological fibrils, as well as, their native interactions. These native contacts represent hydrogen bonds (HB), and hydrophobic and ionic bridges interactions. Moreover, we consider contacts between amino acids in individual chains with sequential distance |i − j| > 4. The parameters *σ*_ij_ are given by *r*_ij0_/2^1/6^, where *r*_ij0_ is the distance between two *C_α_* atoms that form the native contact. The last term in Eq.(1) simply describes the repulsion between non-native contacts. Here, we take *r*_cut_ = 4 Å. Moreover, our terminology for the ‘contacts’ in this manuscript, is as follows: i) intrachain contacts are considered those within a single chain, ii) interchain contacts are between two chains in a side-by-side configuration and iii) the intersheet contacts are found along the symmetry axis (see Fig. 2). Below, we provide details on the different types of mechanical deformation, *i.e*. tensile, shear, and indentation processes.

**Figure 2:**
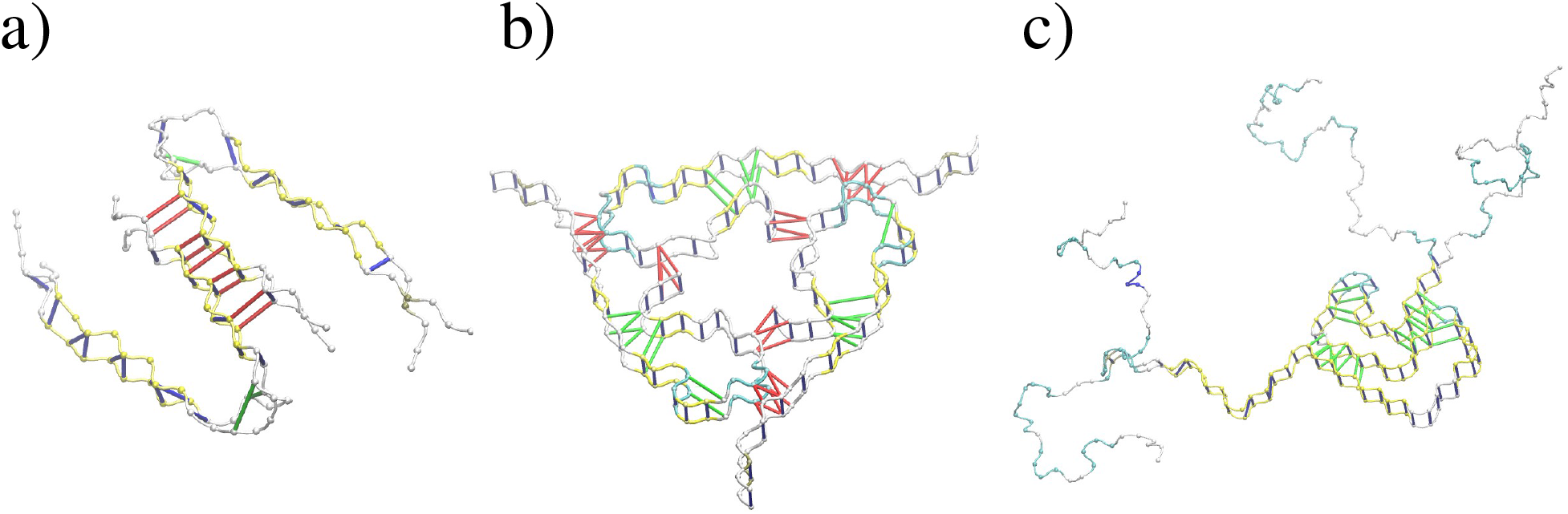
Coarse-grained representation of the biological fibrils presented in Fig.1. We illustrate the three types of ‘native contact’ interactions considered in our study: i) intrachain contacts (green), ii) interchain contacts (red) iii) intersheet contacts (blue).

### Mechanical and thermodynamics characterization through a CG model

In our previous work [36], we have constructed a computational protocol for performing several types of mechanical deformation *in silico* (see Fig. 3). Such processes can be carried out at constant speed or force contact-modes. Here, we explore the former as it provides a dynamic picture of the whole process and it enables the characterisation of the mechanics during the early deformation stages. Moreover, we employ the CG simulation for the validation of the elastic theory. This is done by calculating the coefficient “*n*” in the force *versus h^n^* indentation curves. In particular, we found *n* = 3/2 in the linear regime, which corresponds to the Hertzian theory [12].

**Figure 3:**
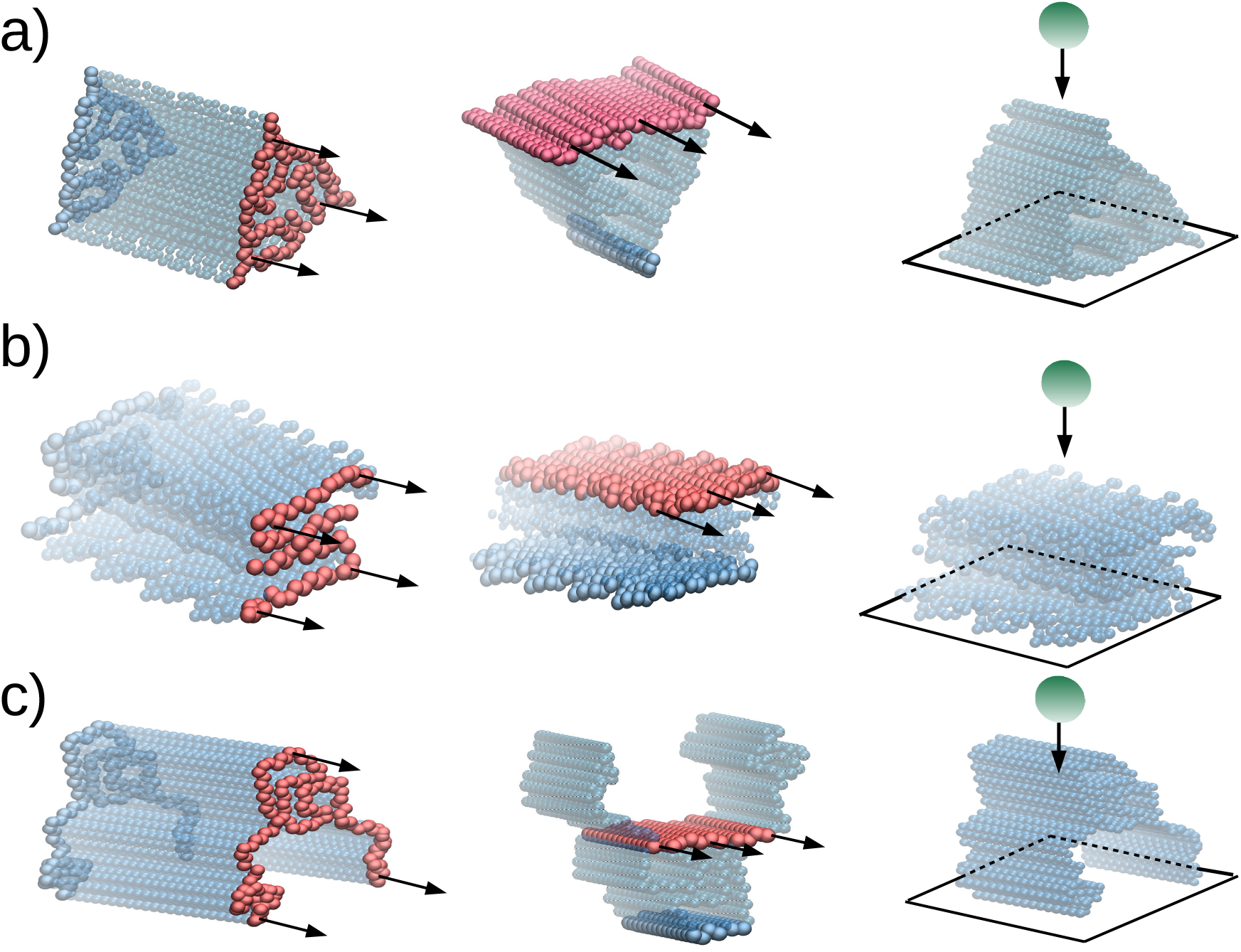
For the cases of Fig. 1, we present schematically each deformation process. Left side shows tensile, middle panel the shearing, and right panel the indentation processes. The set of *C_α_* atoms anchored in each processes are shown in solid blue colour, the ones which are moving at a speed *ν*_pull_ are shown with red colour, and the indenter bead in green. Arrows indicate the direction of pulling. In the case of indentation, a potential 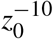 has been used to model the basis plane, where *z*_0_ is the distance between the plane and the CG beads. Top panel shows the structure of *β* A_40_ (PDB id: 2M4J), middle panel for *β* A_40_ (PDB id: 2LMO) and bottom panel *α*-syn (PDB id: 2NA0)

#### Tensile deformation

The exerimental calculation of the stress–strain data in the nanoscale can be done by optical tweezers (OT) [50], AFM base-force spectroscopy [51], or by the design of a sophisticated microelectromechanical systems (MEMS) [52]. These techniques have been successfully used to predict elastic properties of several biomolecules. However, OT are limited to applied loads below 0.1 nN and AFM has delicate calibration issues associated with a systematic deformation of samples with same length. In practice, all-atom simulation does not suffer from any of those drawbacks, but it can not be used in biological systems. Instead, CG models are more suitable to achieve the experimental length and time scales.

In practice, we set harmonic potentials to the furthest bottom and top particles of the protein. Then, we take values for the elastic constants equal to *k*_bottom_ = 100 kcal/mol/Å and *k*_top_ = 0.1 kcal/mol/Å for the top part of the fibril. The top part is moving with pulling speed equal to *ν*_pull_ = 5 × 10^−5^ Å/ns. As a result of tensile deformation, the fibril stretches from a reference length (*L*_0_) to *L*, and the strain is given by *ϕ* = (*L* − *L*_0_)/*L*_0_. The stress is defined by the total force acting on the springs *k*_top_ divided by the cross-sectional area, *A*, of the sample. This area is calculated as follows [53]: for a given set of Cartesian points, it determines the smallest convex polygon containing all the given points. Then, we monitor the elementary area of such polygon during the simulation.[54] From the stress–strain plot one can derive the corresponding tensile Young modulus, *Y*_L_.

#### Shear deformation

The experimental techniques employed before for determination of the *Y_L_* are not transferable for the calculation of the shear modulus (*S*) at the nanoscale. In this respect, an improved version of the single three-point bending technique was developed for the calculation of *S* [55]. It combines a movement along the *z*-axis (perpendicular to the main fibril axis) with a continuous scanning motion along the main fibril axis. In this way, the slope *dF*/*dz* enables a better calculation of the bending modulus (*Y_b_*) and as a result a more accurate value of *S*. In comparison to its predecessor, this technique reduces the error in the value of *S* up to 12% in the case of collagen fibrils [55], but it still relies on the correct estimation of fibril diameter. As above, here CG model helps to devise a protocol where simple shear planes can be applied on a set of atoms and typical response allows in a straightforward manner the calculation of *S*. In this case, we only couple the *C_α_*-atom from the top (*k*_top_) and the bottom (*k*_bottom_) planes. The strain is defined by *ϕ* = *x*/*y*, where *x* is the displacement of the top plane and *y* is the height of the fibril (see Fig. 3). The shear-stress is calculated as the total force acting on the top plane divided by the area of the plane (see in Table 1 the reference *C_α_*-atom used to define the top plane). From the stress–strain relation one can derive the corresponding shear Young modulus, *S*.

**Table 1:** List of geometric parameters of the fibril structures used to determine the *Y*_L_, *Y*_T_, and *S*. Bottom line shows the protein segment used to define the shear plane as illustrated in Fig. 3.

| $A\beta_{40}$ | 2LMO | 2MJ4 | 2MVX |
| --- | --- | --- | --- |
| Initial length, $L_0$ [nm] | $41.10 \pm 0.23$ | $42.21 \pm 0.34$ | $29.10 \pm 0.31$ |
| Cross-section area, $A$ [nm<sup>2</sup>] | $16.02 \pm 0.20$ | $21.11 \pm 0.33$ | $19.20 \pm 0.41$ |
| Shear plane area, $A$ [nm<sup>2</sup>] | $160.01 \pm 0.11$ | $170.20 \pm 0.41$ | $131.00 \pm 0.41$ |
| residue-id involved in shear plane | Gln15–Asp23 | Asp1–Ala23, Asp1' | Gly9–Gly24 |
| $A\beta_{42}$ | 5OQV | 2NAO | |
| Initial length, $L_0$ [nm] | $29.30 \pm 0.23$ | $29.10 \pm 0.31$ | |
| Cross-section area, $A$ [nm<sup>2</sup>] | $17.30 \pm 0.11$ | $14.20 \pm 0.34$ | |
| Shear plane area, $A$ [nm<sup>2</sup>] | $123.00 \pm 0.10$ | $140.10 \pm 0.11$ | |
| residue-id involved in shear plane | Tyr10–Asp23 | Asp1–Asp7, Glu22–Gly25 |  |
| α-syn | 2N0A | | |
| Initial length, $L_0$ [nm] | $45.20 \pm 0.31$ | | |
| Cross-section area, $A$ [nm<sup>2</sup>] | $11.30 \pm 0.41$ | | |
| Shear plane area, $A$ [nm<sup>2</sup>] | $160.00 \pm 0.24$ | | |
| residue-id involved in shear plane | Lys45–Glu105 |  |  |

#### Indentation deformation

One of the empirical techniques used to estimate *Y_T_* modulus is AFM nanoindentation. The wide range of applications of AFM technique span from biomolecules to single cells [31,56,57]. The AFM nanoindentation force–distance curves typically depend on the correct determination of the cantilever stiffness and measurements of biological fibrils located at the center of the fibril are only considered. The former refers to the way that the indentation load is measured by the deflection of the AFM cantilever. The latter is an assumption of the seminfinity half-space approximation. Once the AFM data is obtained, it requires the interpretation by a contact theory. There is not any experiment at the nanoscale where the influence of the indenter could be neglected. Depending on the type of forces between the indenter and the biomaterial, we might describe the process by non-adhesive [12] or adhesive contact theories [58,59]. Here, we suggest our particle-based CG method as a tool to idealize the nanoindentation process. It is worth noting that we prevent any possible adhesion between the indenter and the fibril by placing a divergent interaction between the tip and the *C_α_* atom, and hence other models [58,60] with such features are not considered. moreover we chose the Young modulus of the indenter equal to ∞. Moreover, we define each system in the limit of the Hertzian theory [12]. The indenter is a sphere with a radius of curvature *R*_ind_ that moves towards the fibril with a speed *ν*_ind_. Then, the penetration or indentation depth (*h*) is measured from the first tip–particle interaction (or contact) and the associated indentation force (*F*) is calculated until the indenter stops being in contact with the fibril. From Hertz’s relation, 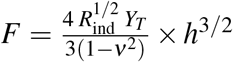, where *ν* is the Poisson coefficient, in this case equal to 0.5. This value corresponds to a homogeneous deformation in the *xy* plane. From Hertz’s equation, we derive the transverse Young modulus, *Y*_T_, in the linear regime of the *F* − *h* curve.

#### Thermodynamic characterization

The study of the thermal stability in the case of A*β* fibrils faces serious difficulties, stemming from the requirement for controlled in *vitro* preparation of samples with well-ordered A*β*_40_ or A*β*_42_ fibrils. In this regard, our CG simulation is an ideal protocol as it enables the calculation of the melting temperatures for well-ordered A*β* fibrils. To assess the thermal stability of the fibril, we compute the probability of finding the protein in the native state, *P*_0_, as a function of the temperature *T*. We define the temperature of thermodynamic stability, *T_m_*, for the case 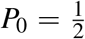. To study the thermodynamic properties of the biological fibrils, we carried out overdamped Langevin dynamics simulations. The simulations were performed for 35 different temperatures, *T*, which were uniformly distributed in the interval 0.1-0.7 *ε*/k_B_. Each simulation was 10^4^*τ* long after running the systems for10^3^*τ* in order to reach equilibrium. In our studies, the unit of time, *τ*, is of the order of 1 ns. For this range of temperatures and time scales, we did not observe any dissociation or unfolding events for the fibrils. The deviation of the fibril structure from its native state was computed by means of the root mean square deviation (RMSD), which is defined as follows:

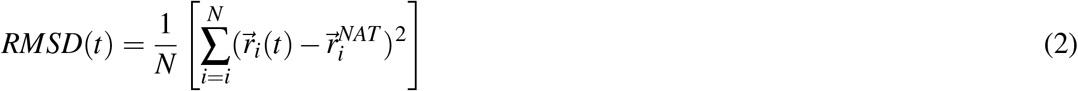

where 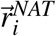 denotes the positions of *C_α_*-atoms in the native state and 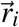 are positions of the *C_α_*-atoms at time t after superimposing the native structure. After equilibration, RMSD fluctuates around an average value, 〈*RMSD*〉, which is a function of temperature *T*. In our case, the observed deviations from the native state in terms of RMSD are small.

## Results and Discussion

### Tensile deformation

Our results for tensile deformation for all studied cases are illustrated in Fig. 4. The initial length (*L*_0_) is measured after an equilibration of 100 *τ*. The cross-section area (A) for each system is monitored during the simulation and is shown as a function of strain in the insets of Fig. 4. The deviations are small compared to the mean value, especially in the case of *β*-amyloid fibrils. Hence, we calculated the stress using the average value of *A*. The values of the cross-section areas and the initial length for each fibril are listed in Table 1. The theoretical values of *Y_L_* have been obtained for *ν*_pull_ = 0.0005 Å/*τ* as listed in Table 2, next to the experimental values for the sake of comparison. In our studies, the deformation is carried out along the main axis of symmetry (see Fig. 1) for *Aβ* and *α*-syn fibrils. We find that the type of *Aβ* fibril plays a more important role in the mechanical properties than the symmetry of each fibril. This becomes apparent by comparing the values of the tensile Young moduli between A*β*_40_ and A*β*_42_. Our discussion is based on the average values of *Y_L_*. In the case of *Aβ*_40_, *Y_L_* = 2.1 GPa, while for A*β*_42_ this value is 2.4 GPa. The value *Y_L_* = 2.3 GPa in the case of *α*-syn seems to be half way between the *Aβ*_40_ and *Aβ*_42_ fibrils. Moreover, our *Y_L_* values are close to the experimental values of collagen fibril equal to 1.9-3.4 GPa [61]. The bottom panels in Fig. 4 illustrate the distributions of lengths for the ‘native contacts’ (intrachain, interchain, and intersheet) as defined in our CG model (Fig. 2). We observe that the intersheet contacts become stretched, an effect that is independent of the system in terms of symmetry or type of individual chains (40 or 42 *β*-amyloid). In contrast, the interchain contacts, which keep together *Aβ* chains in the cross-section area, reduce their length. Moreover, in the case of *α*-syn there are no interchain contacts given that there is only one chain at the cross-section. In this case, only the intrachain contacts stretch during tensile deformation. A similar mechanism is found in *Aβ* fibrils (data not shown), which is consistent with the expectation to maintain the cross-section area constant in the linear regime, used to calculate the Young modulus.

**Figure 4:**
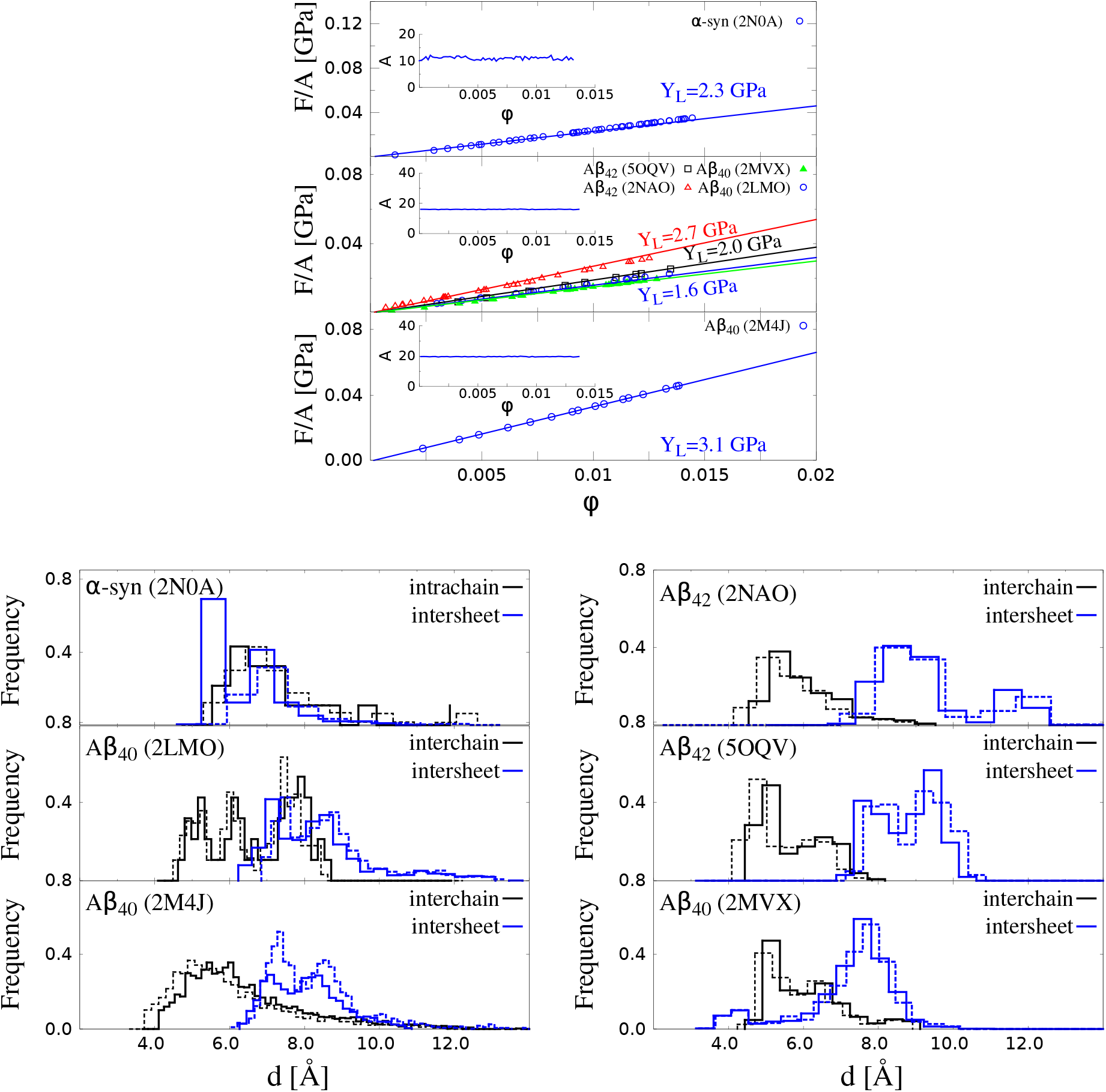
Results on tensile deformation. The top panel shows stress–strain curves of *α*-synuclein, three A*β*_40_ and two A*β*_42_ fibrils. Circles correspond to *ν* = 0.0005Å/*τ*. The error bars are the same as the symbol size and they are based on 50 independent simulations for each structure. The insets show the corresponding cross-section areas in nm^2^ for the corresponding pulling speed. The lower panel shows the distributions of HB lengths for *ϕ* = 0 (solid lines) and for a finite strain *ϕ* corresponding to the end of the linear regime (dashed lines): for *α*-synuclein the final *ϕ* = 0.014, while ϕ = 0.012 for A*β* amyloids.

**Table 2:** The elastic moduli for the A*β*_40_, A*β*_42_ and *α*-syn from experiment and our CG model. in this paper. The structural symmetry of *β*-amyloid (if specified in the literature) is given next to the PDB entries. The experimental results regarding indentation for A*β*_42_ and *α*-syn have been taken from Ref. [30]. The experimental values for the shear modulus (*S*) for *β*-amyloids have been taken from Ref. [62], whereas the experimental value of *S* and *Y*_L_ for *α*-syn are currently unknown.

| Tensile ($Y_L$)/PDB id | Symmetry | $A\beta_{40}$ | $A\beta_{42}$ | α-syn |
| --- | --- | --- | --- | --- |
| 2LMO | 2-fold | $1.6 \pm 0.1$ | | |
| 2MJ4 | 3-fold | $3.1 \pm 0.1$ | | |
| 2MVX | 2-fold | $1.5 \pm 0.1$ | | |
| 5OQV | 2-fold | | $2.0 \pm 0.2$ | |
| 2NAO | 2-fold | | $2.7 \pm 0.2$ | |
| 2N0A | — | | | $2.3 \pm 0.2$ |
| Exp | — | — | — | — |
| Shear ($S$)/PDB id | | | | |
| 2LMO | 2-fold | $0.6 \pm 0.3$ | | |
| 2MJ4 | 3-fold | $1.2 \pm 0.2$ | | |
| 2MVX | 2-fold | $0.4 \pm 0.1$ | | |
| 5OQV | 2-fold | | $1.3 \pm 0.2$ | |
| 2NAO | 2-fold | | $1.8 \pm 0.1$ | |
| 2N0A | — | | | $0.7 \pm 0.2$ |
| Exp | — | $0.1 \pm 0.02$ | — | — |
| Indentation ($Y_T$)/PDB id | | | | |
| 2LMO | 2-fold | $3.0 \pm 0.1$ | | |
| 2MJ4 | 3-fold | $6.0 \pm 0.2$ | | |
| 2MVX | 2-fold | $5.0 \pm 0.1$ | | |
| 5OQV | 2-fold | | $7.0 \pm 0.3$ | |
| 2NAO | 2-fold | | $16.0 \pm 0.4$ | |
| 2N0A | — | | | $13.0 \pm 0.1$ |
| Exp | — | — | $3.2 \pm 0.8$ | $2.2 \pm 0.6$ |

### Shearing deformation

Our results for all systems are presented in Fig. 5. The shear deformation for *Aβ* and *α*-syn fibrils takes place along the same directions as in the case of tensile deformation (see Fig. 3). The initial values of the top-plane areas for each fibril are listed in Table 1. The insets in the left panels of Fig. 5 demonstrate that the area A does not change when shear is applied. The values of shear modulus (*S*) computed for *ν*_pull_ = 0.0005 Å/*τ* are listed in Table 2. In our studies, these values show a large dependence on the type of *Aβ* fibril. We find that *S* for A*β*_42_ is about 1.6 GPa, while for *Aβ*_40_ it is equal to 0.7 GPA. The 2.3-fold increase supports the picture that the *Aβ*_42_ fibril is mechanically more stable than the *Aβ*_40_ [27]. The *S* value for *α*-synuclein is comparable to the *Aβ*_40_. No experimental data on *S* for *α*-synuclein fibril has been reported, but it is expected to comprise the range between 1.4-300 MPa. Both limits are typical of microtubules [63] and collagen [55] systems, which are assemblies of proteins. Main discrepancies between our computational studies and experimental results are expected. One of the sources of divergence is associated with the crystal-like regions, which are present in the biological fibrils during each deformation in silico. The initial structure of fibrils are very close to the minimum free energy state (native). Here, the number of hydrogen bonds that participate in the deformation as a whole is larger as reported by all-atom [4,5]. In contrast, during in vitro self-assembly of neurodegenerative fibrils the fibrilization process is dominated by extended regions of amorphous aggregates. Such regions will induce the overall softening of the fibril and therefore the drop in the elastic modulus.

**Figure 5:**
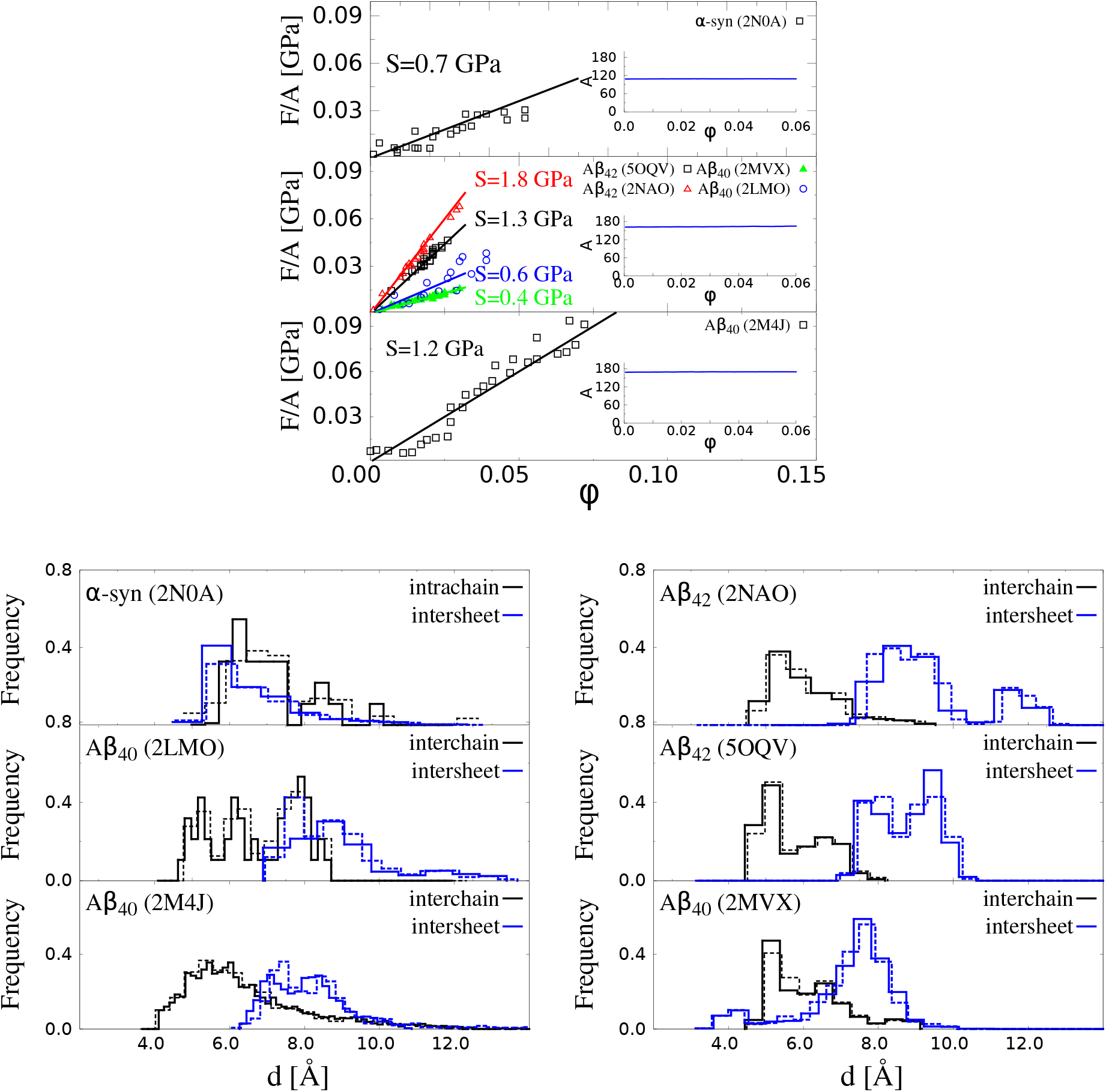
Results for shear deformation. The top panel shows stress-strain curves of *α*-syn and three A*β*_40_ and two A*β*_42_ fibrils. Circles refer to *ν* = 0.0005Á/*τ*. The error bars are the same as the symbol size and they are based on 50 independent simulations for each structure. The inset shows the corresponding cross-section area in nm^2^. The lower panel presents the distributions of the HB lengths for *ϕ* = 0 (solid lines) and for a finite *ϕ* corresponding to the end of the linear regime (dashed lines), which is 0.04 for *α*-syn and 0.025 for A*β* amyloids. Only A*β*_40_ with PDB id: 2M4J has been calculated at strain *ϕ* = 0.05.

The bottom panels in Fig. 5 show the distributions of the characteristic native distances (see Fig. 2 for their definition). For *β*-amyloid and *α*-synuclein fibrils, the intersheet contacts become slightly stretched, but the distances in the interchain contacts within each sheet are not affected in the case of amyloids. The same analogy can be seen for the intrachain contacts in *α*-synuclein. This effect helps the system to keep constant the thickness of the fibril, a condition for the calculation of shear modulus in the linear regime.

### Indentation deformation

Our results for all systems are presented in Fig. 6. The indentational deformation for *Aβ* and *α*-syn fibrils takes place in the normal direction the plane, *z* = 0 and at the position *L* = 1/2*L*_0_ (see Fig. 3).

**Figure 6:**
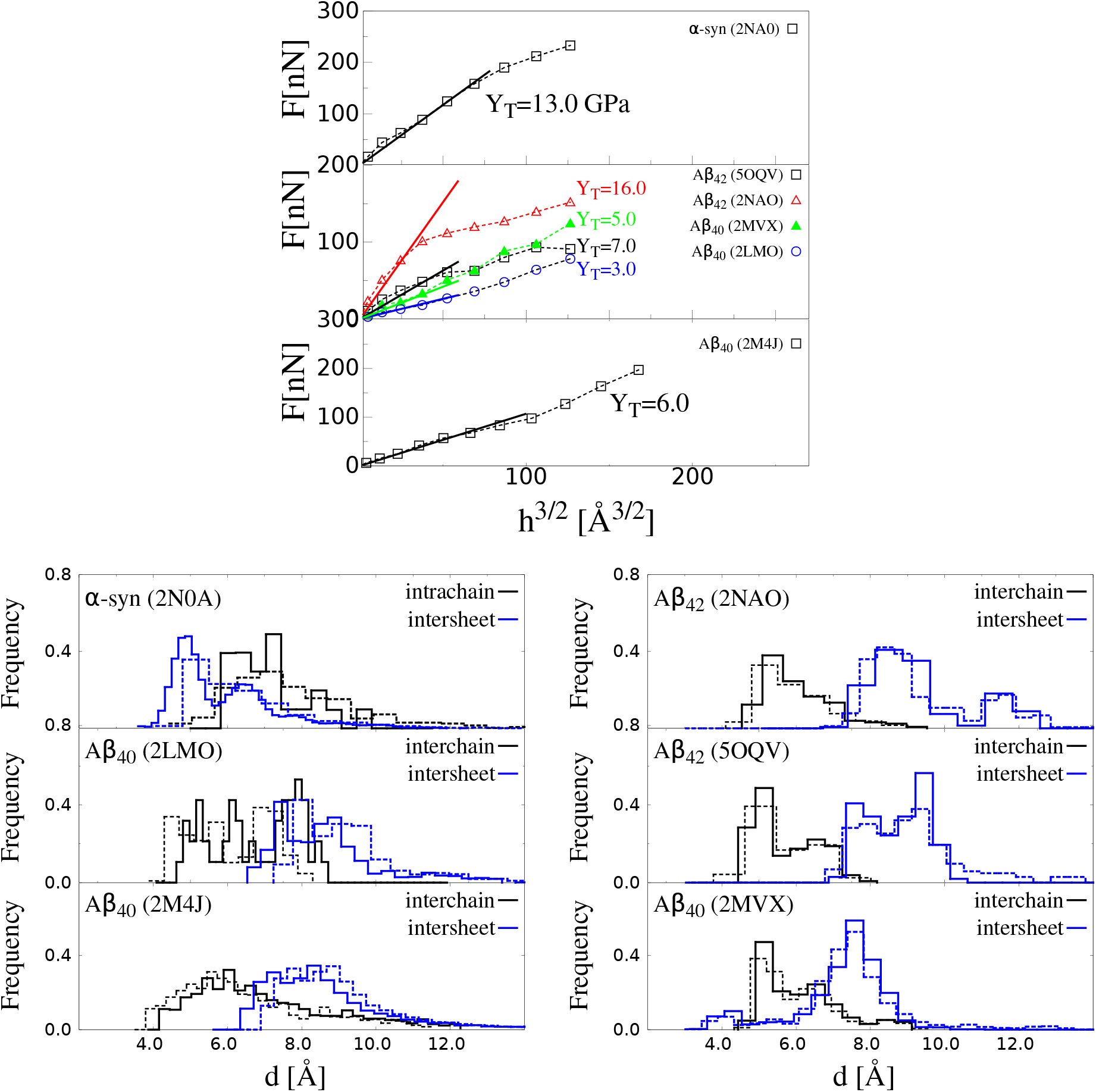
Nanoindentation deformation results for different biological fibrils. The top panel shows plots of force *versus* indentation depth (*h*) for *α*-syn, three A*β*_40_, and two A*β*_42_ fibrils. Square symbols refer to *ν*_ind_ = 0.005 Å/*τ* and *R*_ind_ = 10 nm. The error bars are the same as the symbol size and they are based on 50 independent simulations for each system. The distributions are calculated for *h* = 0 (solid line) and *h* = 20 Å in the case of *α*-syn fibril and A*β* fibrils (dashed lines). Only in the case of *Aβ*_42_ with PDB id: 2NAO the value *h* = 9 Å was considered.

The initial values of the fibril length for each fibril are listed in Table 1. The values of transversal Young modulus (*Y_T_*) computed for *ν*_pull_ = 0.005 Å/*τ* are listed in Table 2. In our studies, for the case of A*β* our results show a large dependence on the type of *Aβ* fibril. We determine that *Y_T_* for A*β*_42_ is about 12 GPa, while for *Aβ*_40_ it is equal to 5 GPA. The 2.5-fold increase supports the picture that the *Aβ*_42_ fibril is mechanically more stable than the *Aβ*_40_ [27]. Because *Aβ*_42_ aggregates faster than *Aβ*40 [64] our findings support the correlation between mechanical stability and aggregation propensity as in ref. [27]. The *Y_T_* value for *α*-synuclein is comparable to the *Aβ*_42_. The experimental data on *Y_T_* for *α*-syn fibril has been reported [30] and it is a factor 2 smaller than A*β*_40_. Such difference is attributed uncontrollable growth of amorphous aggregates during fibrillization that makes softer the fibril. But it is worth mentioning that our theoretical values can be considered as an upper bound and it derived such parameter in the case of highly ordered fibrils. Moreover, the same result has been observed in all-atom simulations studies [5].

The bottom panels in Fig. 6 show the distributions of the characteristic native distances (see Fig. 2 for their definition). For A*β* and *α*-syn fibrils, the intersheet contacts become stretched, but the distances in the interchain contacts within each sheet are shortened in the case of amyloids. The same analogy can be seen for the intrachain contacts in *α*-synuclein.

### Thermodynamic characterization of fibrils

Our results regarding the effect of the temperature for each fibril structure are presented in Fig. 7. We first study the *P*_0_ for all fibrils as a function of the temperature. Fig. 7 (top panel) shows that the probability *P*_0_ of finding the fibrils in the native state is larger for the *Aβ*_40_ and *Aβ*_42_ when compared to *α*-syn at any given temperature. This result is in agreement with a differential calorimetry experiment where it is observed that *T_m_* for *β*-amyloid fibrils is larger than *α*-syn fibrils [65,66]. In terms of the single fibril the *Aβ*_40_ (PDB id: 2MVX) with two-fold symmetry is the most stable at higher temperature (thermophilic character) among the other two-fold and three-fold *β*-amyloids. The calibration of our room temperature is 0.35 *ε*/*k*_B_. In particular, the folding temperature (*T*_f_) defined in our CG model at *P*_0_ equal to 0.5 gives *T*_f_ equal to 0.38, 0.42, 0.44, 0.46, and 0.48 in units of *ε*/*k*_B_ for the amyloids with PDB entry 2LMO, 2MJ4, 2NAO, 5OQV, and 2MVX, respectively. With our calibration of ε, the difference between the most (PDB id: 2MVX) and less (PDB id: 2LMO) thermophilic fibrils is of the order of 85°C. Our results indicate that the *α*-syn fibril is less thermally stable in comparison with the *Aβ* system and this behaviour seems to be intrinsically associated with the extended disordered N-terminus and C-terminus domains. In our model, for *α*-syn we have determined that *T*_f_ is 0.33 *ε*/*k*_B_. The difference in temperature with respect to A*β* with PDB ids 2LMO and 2MVX is 43 °C and 128 °C, respectively. This implies a higher thermodynamic stability of the A*β* systems in comparison with *α*-syn, which may explain the easier formation of *Aβ* fibrils over *α*-syn. Fig. 7 (right side) shows that 〈RMSD〉 is larger in the case of *α*-syn than in the case of *Aβ* fibrils, at any given *T*. In addition, Fig. 7 (bottom panel) presents the RMSF results for all fibrils. We observe that the disordered domains (N- and C-terminus) in *α*-syn are very flexible in comparison with *Aβ* fibrils.

**Figure 7:**
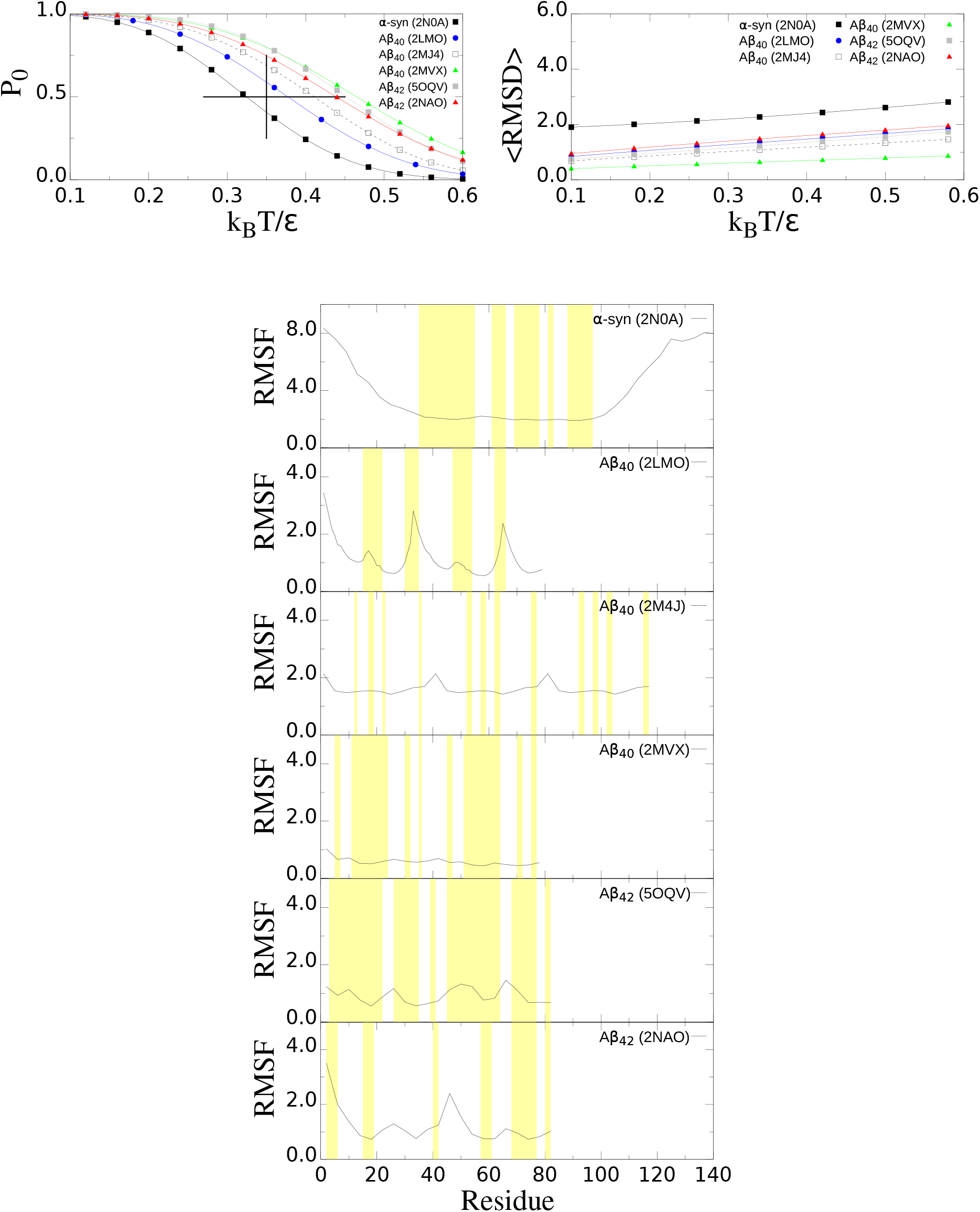
Thermodynamic properties of biological fibrils. Top (left) panel shows the probability of finding the fibrils in the native “ensemble” state, P_0_, as a function of the temperature. The vertical line indicates the room temperature equal to 0.35 *ε*/*k*_B_ and the horizontal line the range of temperatures that offer thermodynamic stability in our model. Top (right) panel illustrates the RMSD of the fibrils. Bottom panel illustrates the root-mean-square-fluctuation (RMSF) at room temperature. The *β*-strand segments in each system are highlighted in yellow.

## Conclusion

We have carried out molecular dynamics simulations to study the elastic properties of two families of biological fibrils, namely, the *β*-amyloid and *α*-syn. The elastic properties of this study are the tensile, shear, and indentation deformations. Overall, our results are in agreement with the corresponding experimental values that could be obtained from the literature. Moreover, our method is sensitive to variations in the chain length and the symmetry of the *β*-amyloid fibril. Our results indicate a higher mechanostability in the case of *β*A_42_ fibrils than in the case of *β*A_40_, namely, 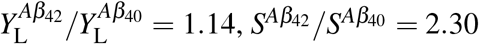, *S*^*Aβ*_42_^/*S*^*Aβ*_40_^ = 2.30, and 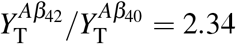 This result is consistent with the results obtained by means of the rupture force [27]. Most importantly, given that the aggregation rate depends on the mechanical stability of the fibrils [27] our study could provide also hints for self-assembly *β*-amyloid and *α*-syn chains. Our results also indicate an elastic anisotropy namely, *Y_T_* > *Y_L_*, for all systems. In the case of *α*-syn fibrils such anisotropy, which is expressed by the difference between *Y_T_* and *Y_L_*, which is almost one order of magnitude. In contrast, in the case of *β*-amyloid fibrils the anisotropy is considerably smaller.

We find that this effect is due to the deformation of the hydrophobic core (segments 61–95). We have also confirmed that the large anisotropy in the case of *α*-syn neither depends on the N-terminus nor the C-terminus domains. Although the the mechanical properties indicate some similar behaviour between *α*-syn and *β*-amyloid fibrils, thermodynamic properties reveal a different behaviour, that is *β*-amyloid fibrils are thermally more stable than *α*-syn fibrils. Hence, *β*-amyloid fibrils are in general more stable at higher temperatures than at room temperature, for example, whereas the opposite effect takes place in the case of *α*-syn fibrils. In this regard, our method can be used to explore systematically the temperature dependence of the mechanical properties (thermoelastic) in biological fibrils at experimental length and time scales.

## Acknowledgements

We thank Claudio Perego for critically reading the manuscript. This research has been supported by the National Science Centre, Poland, under grant No. 2015/19/P/ST3/03541, 2015/19/B/ST4/02721, and No. 2017/26/D/NZ1/00466. This project has received funding from the European Union’s Horizon 2020 research and innovation programme under the Marie Sklodowska-Curie grant agreement No. 665778. This research was supported in part by PLGrid Infrastructure.

